# Emergence Risks from Within? Metagenomic Analysis of Mosquito Viromes from Two Zoos Reveals Mosquito-associated Orthobunyaviruses in the UK

**DOI:** 10.1101/2025.05.01.651672

**Authors:** Jack Pilgrim, Nicola Seechurn, Edward Cunningham-Oakes, Phillipa Dobbs, Javier Lopez, Alain Kohl, Grant L Hughes, Marcus SC Blagrove, Matthew Baylis, Alistair C Darby

## Abstract

In regions naïve to mosquito-borne threats, risk assessments often prioritise incursion events, particularly those driven by climate change and host movement. However, these same factors may also facilitate the emergence of viruses already circulating in local vector populations. The use of metagenomics to assess mosquito viromes has expanded in recent decades, enabling the detection of both arboviruses and insect-specific viruses (ISVs), and helping to fill gaps in our understanding of virus circulation. In particular, zoological collections are potential interfaces for virus transmission between mosquitoes, animals, and humans, making them valuable sentinel sites for surveillance. Although mosquito virome studies have been conducted globally, UK mosquito populations remain comparatively understudied. To address this gap, we investigated the viromes of two key mosquito species, *Culex pipiens* s.l. and *Culiseta annulata*, through metagenomic sequencing of specimens collected at two UK zoos between 2021 and 2022. A total of 4,042 mosquitoes underwent sequencing across 44 pools, leading to the identification of 26 viral genomes, including nine novel species spanning RNA and DNA virus families. Viral distribution patterns revealed viruses shared across all pools, supporting the possibility of a conserved virome between mosquito species. Notably, two novel *Orthobunyavirus* species (Family *Peribunyaviridae*), Atherstone and Deva viruses, were detected. These represent the first documented mosquito-associated orthobunyaviruses in the UK, a genus that includes several known pathogens. Classification using the machine-learning tool MosViR suggests Atherstone virus, present at both zoos, is a putative arbovirus warranting further investigation. Supporting this is the presence of a non-structural protein S (NSs), a known virulence factor in orthobunyaviruses. In contrast, Deva virus had no NSs ORF and was not predicted to be an arbovirus. These findings improve our understanding of UK mosquito viruses, indicate possible risks to people or animals from orthobunyaviruses, and highlight the role of metagenomic surveillance in detecting emerging viral threats.

## Introduction

While malaria transmission persisted in the UK until the early 20th century [1], mosquitoes have since posed limited concern for public or veterinary health due to the absence of other major mosquito-borne diseases. However, the recent emergence of the flavivirus Usutu virus marks the first recorded instance of an established mosquito-borne zoonotic virus in the country [2,3], underscoring the need to better understand mosquito ecology in this region. Additionally, the recent encroachment of the related West Nile virus into northern Europe [4], along with the detection of neglected arboviruses in the region [5], has heightened concerns about the potential for UK mosquitoes to transmit emerging arboviruses [6,7]. Importantly, the emergence of arboviruses such as Schmallenberg virus in Northern Europe [8] highlights that local vector species, given suitable conditions, can support the emergence of novel pathogens. This underscores the need for proactive surveillance to better characterise the UK’s vector populations and anticipate future threats.

Arbovirus emergence can result both from the incursion of exotic viruses [9,10] and from the amplification of endemic viruses already present in local vector populations [11,12]. Factors such as climate change, host movement, and environmental modification are known to facilitate both processes, influencing vector distribution, abundance, and virus transmission dynamics [13,14]. Zoological collections represent unique ecological interfaces where local vectors, exotic and native wildlife, and humans coexist in close proximity. Such settings are likely to harbour high densities of mosquitoes, attracted by a steady availability of vertebrate hosts and suitable breeding habitats. In addition, the importation of non-native animal species raises the possibility of exotic viruses being introduced and maintained within these environments [15]. Simultaneously, zoos may also act as sentinel sites for detecting endemic or under-recognised pathogens [16,17], particularly those circulating silently in native mosquito populations [18].

Innovations in sequencing technologies over the past decade have improved scalability, affordability, and the detection of both known and novel viruses in mosquito populations [19]. Metagenomic techniques have expanded our understanding of mosquito viromes by uncovering both arboviruses and a wide array of insect-specific viruses (ISVs) [5,20–25]. These approaches also enhance surveillance by detecting known arboviruses in previously unsurveyed regions and flagging novel viruses with arboviral potential [26–28]. As these approaches are agnostic to prior sequence knowledge, they can reveal a broader range of viruses than traditional arbovirus surveillance methods reliant on targeted PCR or serological inference. More recently, large-scale mosquito transcriptomic datasets, combined with machine learning, have shown promise for identifying putative arboviruses for targeted investigation, offering a powerful adjunct to existing surveillance frameworks [29].

While, the detection of arboviruses is crucial for anticipating and preventing outbreaks, simultaneously unveiling ISVs also has significant implications, as they can influence vectorial capacity [30,31]. ISV arbovirus-modulating impacts are being documented more frequently. For instance, cell-fusing agent virus (CFAV) has been shown to decrease the dissemination of dengue and Zika viruses in *Aedes aegypti* [32], while *Culex* flavivirus modulates the capacity of *Cx. pipiens* s.l. to transmit WNV under laboratory conditions [33]. These effects may be explained by superinfection exclusion, a phenomenon in which the presence of an ISV limits or prevents subsequent infection of the same cell by an arbovirus [34–37]. Alternatively, ISV-mediated activation of immune pathways such as Jak/Stat, Toll, or IMD may influence arbovirus replication or transmission within mosquito populations, potentially offering strategies for arbovirus control. [30]. Furthermore, certain DNA ISVs have been shown to reduce mosquito longevity and, coupled with their capacity to be transmitted either vertically or horizontally, make them potential agents for mosquito biocontrol [38,39]. Beyond their impact on vectorial capacity, ISVs can also shed light on the evolutionary transition from insect-specific to dual-host viruses, as many extant arboviruses are believed to have originated from insect-restricted ancestors [40].

This study aimed to detect and characterise the diversity of arboviruses and ISVs present in mosquito populations sampled from two UK zoos. To this end, a metagenomic analysis of UK *Culex pipiens* s.l. and *Culiseta annulata*, insect vectors implicated in the transmission USUV and WNV [41,42], was undertaken. Identification of viruses present within these populations will enhance preparedness and responsiveness to emerging arboviral threats by guiding vector surveillance, population control strategies, and efforts to modulate mosquito vector competence.

## Methods

### Specimen collections

Mosquito specimens were collected and morphologically identified following the methods described in Seechurn *et al.* 2024 [43]. Briefly, adult mosquitoes were trapped using multiple trap types at Chester Zoo and Twycross Zoo during the spring and summer of 2021 and 2022. Mosquitoes were stored individually in Eppendorf tubes at -80°C before being pooled by zoo, species, sex and year. In total, 3,970 *Cx. pipiens* s.l. mosquitoes were grouped into 41 pools, with an average of 94.4 individuals per pool (Table 1). Additionally, three pools of *Culiseta annulata* mosquitoes were processed: two from Chester Zoo (n = 48 and n = 60) and one from Twycross Zoo (n = 65).

**Table 1.** Information on mosquito pools collected from Chester and Twycross Zoos, including species, sex, location, collection year and pool size. Cx. pip = *Culex pipiens*, Cs. ann = *Culiseta annulata*

| Sample pool | Mosquito species | Sex | Zoo location | Year | Number pooled |
| --- | --- | --- | --- | --- | --- |
| NS1 | *Cx. pip* | F | Chester | 2021 | 100 |
| NS2 | *Cx. pip* | F | Chester | 2021 | 100 |
| NS3 | *Cx. pip* | F | Chester | 2021 | 100 |
| NS4 | *Cx. pip* | F | Chester | 2021 | 100 |
| NS5 | *Cx. pip* | F | Chester | 2021 | 93 |
| NS9 | *Cx. pip* | F | Chester | 2021 | 97 |
| NS10 | *Cx. pip* | F | Chester | 2021 | 96 |
| NS11 | *Cx. pip* | F | Chester | 2021 | 98 |
| NS12 | *Cx. pip* | F | Chester | 2021 | 97 |
| NS14 | *Cx. pip* | F | Chester | 2021 | 40 |
| NS15 | *Cx. pip* | F | Twycross | 2021 | 100 |
| NS16 | *Cx. pip* | F | Twycross | 2021 | 100 |
| NS17 | *Cx. pip* | F | Twycross | 2021 | 98 |
| NS19 | *Cx. pip* | F | Twycross | 2021 | 100 |
| NS18 | *Cx. pip* | F | Twycross | 2021 | 113 |
| NS20 | *Cx. pip* | F | Twycross | 2021 | 100 |
| NS21 | Cx. pip | F | Twycross | 2021 | 100 |
| NS22 | Cx. pip | F | Twycross | 2021 | 100 |
| NS23 | Cx. pip | F | Twycross | 2021 | 100 |
| NS24 | Cx. pip | F | Twycross | 2021 | 95 |
| NS25 | Cx. pip | F | Twycross | 2021 | 100 |
| NS26 | Cx. pip | F | Twycross | 2021 | 99 |
| NS29 | Cx. pip | F | Chester | 2022 | 93 |
| NS30 | Cx. pip | F | Chester | 2022 | 100 |
| NS30A | Cx. pip | F | Chester | 2022 | 98 |
| NS31 | Cx. pip | F | Chester | 2022 | 100 |
| NS32 | Cx. pip | F | Chester | 2022 | 100 |
| NS33 | Cx. pip | F | Chester | 2022 | 97 |
| NS34 | Cx. pip | F | Chester | 2022 | 100 |
| NS35 | Cx. pip | F | Chester | 2022 | 101 |
| NS36 | Cx. pip | F | Chester | 2022 | 97 |
| NS37 | Cx. pip | F | Chester | 2022 | 103 |
| NS38 | Cx. pip | F | Chester | 2022 | 59 |
| NS40 | Cx. pip | F | Twycross | 2022 | 105 |
| NS41 | Cx. pip | F | Twycross | 2022 | 100 |
| NS42 | Cx. pip | F | Twycross | 2022 | 98 |
| NS43 | Cx. pip | F | Twycross | 2022 | 98 |
| NS46 | Cx. pip | F | Twycross | 2022 | 98 |
| NS47 | Cx. pip | F | Twycross | 2022 | 98 |
| NS48 | Cs. ann | F | Chester | 2021 &<br>2022 | 48 |
| NS49 | Cs. ann | F | Chester | 2022 | 60 |
| NS50 | Cs. ann | F | Twycross | 2021 &<br>2022 | 65 |
| NSM1 | Cx. pip | M | Chester | 2021 &<br>2022 | 74 |
| NSM3 | Cx. pip | M | Twycross | 2021 &<br>2022 | 24 |

### Sample preparation

Pools were placed in ZR BashingBead Lysis Tubes (Zymo Research) with 600 µl of phosphate-buffered saline (PBS). Samples were homogenized using the FastPrep-24™ Classic at 5 m/s for 40 seconds. The homogenates were rested for 5 minutes, then centrifuged at 16,000 x g for 5 minutes at 4°C. A total of 400 µl of supernatant was filtered through a 5 µm filter with an omniphobic polyvinylidene fluoride (PVDF) membrane (Millex-SV, Merck). Next, 300 µl of filtrate was passed through a 0.8 µm polyethersulfone (PES) membrane filter (Sartorius Vivaclear Mini), followed by a final filtration through a 0.45 µm PES membrane filter (Corning Costar Spin). Up to 130 µl of the remaining filtrate was incubated at 37°C for 1.5 hours with the following reagents per reaction: 3 µl Baseline Zero (1 U/µl, Cambio), 7 µl TURBO DNase (2 U/µl, ThermoFisher, Ambion), 2 µl RNase ONE Ribonuclease (10 U/µl, Promega), and 8 µl 10X DNase buffer (ThermoFisher, Ambion). Total RNA was extracted from the incubated filtrate using the QIAamp Viral RNA Mini Kit (Qiagen) without carrier RNA, following the manufacturer’s instructions.

### Sequence-independent single primer amplification (SISPA)

Sequence-independent single primer amplification (SISPA) was performed as described by Greninger *et al.* [44], with minor modifications. Briefly, 5 µL of total nucleic acid (TNA) was incubated at 65°C for 5 minutes with 40 pmol of the primer Sol-A (5’— GTT TCC CAC TGG AGG ATA NNN NNN NNN—3’) and 1 µL of 10 mM dNTP mix. Reverse transcription was then carried out using SuperScript IV Reverse Transcriptase (Thermo Fisher Scientific) according to the manufacturer’s protocol, with the addition of the ribonuclease inhibitor RNaseOUT (Life Technologies). Second-strand synthesis was performed using Sequenase Version 2.0 (Thermo Fisher Scientific). Reaction mixtures were cleaned using Agencourt AMPure beads following the AMPure XP protocol.

Each reaction was then prepared to a final volume of 50 µL, consisting of 5 µL of cDNA, 25 µL of Q5 master mix (NEB), 1 µL of SolB primer (5’— GTT TCC CAC TGG AGG ATA—3’) at 100 pmol/µL, and 19 µL of nuclease-free water. PCR cycling conditions were as follows: initial denaturation at 98°C for 30 seconds, followed by 35 cycles of 98°C for 10 seconds, 54°C for 30 seconds, and 72°C for 1 minute, with a final extension at 72°C for 10 minutes. Agencourt AMPure bead clean-up was performed to purify PCR products.

The cDNA concentration of each sample was assessed using a Qubit™ Flex Fluorometer and the Qubit™ dsDNA High Sensitivity kit (Thermo Fisher). PCR products were visualized with an Agilent 2100 Bioanalyzer, and cDNA purity (260/280 and 260/230 ratios) was measured using a NanoDrop™ 1000 spectrophotometer.

### Library preparation

cDNA fragmentation was performed using the NEBNext® Ultra™ II FS DNA Library Prep Kit for Illumina (New England Biolabs) following the manufacturer’s instructions. Samples were cleaned using Agencourt AMPure beads. Indexing was completed with NEBNext Multiplex Oligos for Illumina, followed by two additional Agencourt AMPure bead clean-ups. Samples were eluted in nuclease-free water, and quality control was conducted as described above. Samples were stored at -20°C until sequencing, which was performed using the Illumina NovaSeq 6000 System with 150 bp paired-end reads by the Centre of Genomic Research (CGR), University of Liverpool.

### Bioinformatics

FASTQ files were trimmed using Cutadapt version 1.2.1 [45] to remove adapter sequences. The quality of forward and reverse reads was assessed and visualized with FastQC version 0.12.1 [46]. Kraken2 v2.1.2 [47] was used for taxonomic profiling of raw reads to provide preliminary data on viral species abundance within the dataset, utilizing paired-end reads and the viral RefSeq database database for comparisons.

*De novo* assembly was performed using SPAdes v3.15.5 [48] with standard k-mer sizes and the *metaspades* flag with resulting contigs over 500 nucleotides filtered for further processing using SeqKit v0.10.0 [49]. A local amino acid BLAST database was generated from the Virus-Host DB [50] to facilitate the identification of putative viral contigs. To this end, BLASTx (blast+ v.2.15.0) was used with an E-value threshold of 1 x e^-5^ and a minimum query coverage per high-scoring segment (HSP) of 30% (-qcov_hsp_perc 30). As viral metagenome-assembled genomes (MAGs) are often fragmented due to inter-genome shared regions, within-population variants and intra-genome repeats, assemblers often create breaking points in de Bruijin graphs. Therefore, contigs with the same BLAST hit and similar depth of coverage were mapped back to the closest BLASTn hit from NCBI and manually joined to create a scaffold. The quality and completeness of the final viral contigs were assessed using CheckV v0.7.0 [51], with only genomes classified as at least medium-quality (≥50% completeness) included in further analysis. The validity of reconnected breakpoints was verified by mapping reads back to the final contigs using bwa-mem2 v2.2.1 [52], followed by visualization in IGV v2.12.3 [53]to confirm accurate genome reconstruction.

### Phylogenetics and taxonomic assignment

Final viral contigs which passed quality control and had at least one complete gene designated by ORFfinder (https://www.ncbi.nlm.nih.gov/orffinder/), which identifies complete ORFs (open-reading frames), were then taken forward for taxonomic assignment. To achieve this for RNA viruses, RNA-dependent RNA polymerase (RdRp) genes from contigs were subjected to phylogenetic tree reconstruction. For DNA viruses, either the replication-associated protein (Rep) or NS1 protein were used for analysis. The analysis included the closest BLASTp hits to contigs, along with exemplar viral species from closely related genera, as designated by the International Committee on Taxonomy of Viruses (ICTV). MAFFT v7.4 [54] aligned genomes with the *E-INSI* algorithm before using IQtree v1.6.12 [55] to construct maximum likelihood trees with 1000 ultrafast bootstrap replicates and default parameters. Contigs that clustered in a monophyletic group within a recognised genus or family were accordingly assigned to that corresponding taxonomic level. Furthermore, novel viruses were classified based on their fulfilment of the ICTV-designated criteria for a novel species. In cases where no designated species were available for a given genus, a threshold of <90% amino acid identity for the RdRp protein was applied relative to the nearest NCBI match.

To assess the likelihood of a viral genome being mosquito-associated or a putative arbovirus, specific criteria were applied. A virus was considered mosquito-associated if it fulfilled at least two of the following criteria: **(i)** it clustered phylogenetically with other mosquito-derived viruses, **(ii)** it clustered with known arboviruses, or **(iii)** it was detected in multiple mosquito pools. A virus was designated as a putative arbovirus if it **(i)** clustered within a clade containing known arboviruses, and **(ii)** received a probability score >0.60 from the MosViR tool, which applies machine learning models to predict arboviral potential based on genomic features [29]. For orthobunyaviruses, further support for arboviral potential was provided by the detection of a non-structural protein S (NSs) open reading frame (ORF) in the S segment, given its known role in immune evasion and maintenance of infection in vertebrate hosts [56,57]. NSs presence was determined by identifying ORFs located on the same strand as the nucleocapsid (N) gene, within 100 nucleotides of the N gene start codon, and encoding ≥65 amino acids. The presence or absence of NSs ORFs was included in phylogenetic annotations, alongside the Centers for Disease Control and Prevention (CDC) Arbovirus Catalog (ArboCat) classification of known orthobunyaviruses [58], to assist in interpreting arbovirus risk.

### Diversity and distribution of viruses

A viral abundance table was then generated using the methods described by De Coninck et al [25]. Briefly, bwa-mem2 was used to map reads back to the final contigs, and CoverM v0.7.0 [59] was then employed to determine a contig’s presence in a pool, considering it present if at least 50% of its length was covered. The resulting total viral read count for each pool were then converted into a matrix and visualized using the pheatmap package in R [60]. In addition, the relative abundance of viral families in each pool was visualized using ggplot2 [61].

To investigate genomic diversity between viral genomes across pooled samples, Average Nucleotide Identity (ANI) was calculated. BAM files generated from bwa-mem2 mapping were used as input for samtools v1.21 [62] using the *mpileup* flag to generate pileup files. Consensus sequences were then constructed using iVar v1.4.3 [63], with only genomes meeting a minimum of 10× depth of coverage and at least 90% breadth of coverage retained for further analysis. ANI values were computed using FastANI v1.34 [64], and the resulting matrices were visualized using pheatmap.

## Results

A total of 632.8 million reads were generated from the pooled mosquito samples (mean = 14.4 million reads per sample), with 252.6 million reads identified as viral (mean = 5.7 million reads per sample) (Figure 1C). Subsequent assembly and quality control (Supplementary Table S1) revealed the (near-) complete genomes of 21 RNA viruses and 5 DNA viruses, 9 of which were classified as novel (Table 2).

**Figure 1.**
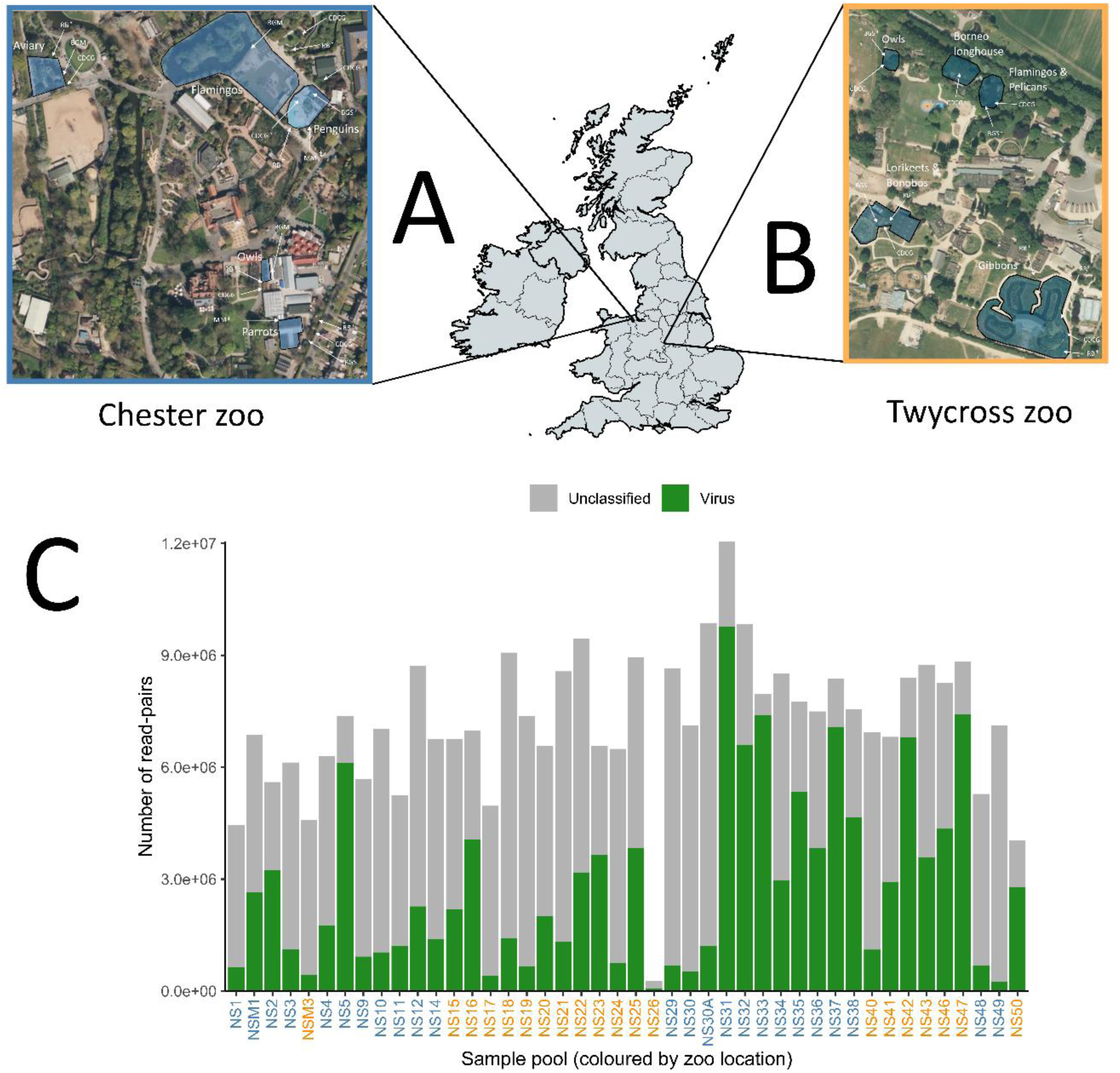
Location of trapping sites from 2021 to 2022 at two zoos: **(A)** Chester Zoo and **(B)**. Blue-shaded regions indicate enclosures where traps were placed. †: Traps only present in 2021. ‡: Traps only present in 2022. BGM: Biogents-Mosquitaire trap. BGS: Biogents Sentinel-2 trap. MM: Mosquito Magnet trap. CDCG: CDC Gravid trap. RB: Resting box trap. **(C)** The number of viral reads detected in each pooled sample, as identified by Kraken2. NS1-NS47 correspond to female *Culex pipiens* s.l., NS48-NS50 correspond to female *Culiseta annulata*, and NSM1 and NSM3 correspond to male *Culex pipiens* s.l. Orange labels = Twycross zoo. Blue labels = Chester zoo. Background satellite imagery courtesy of OpenStreetMap contributors.

**Table 2.**
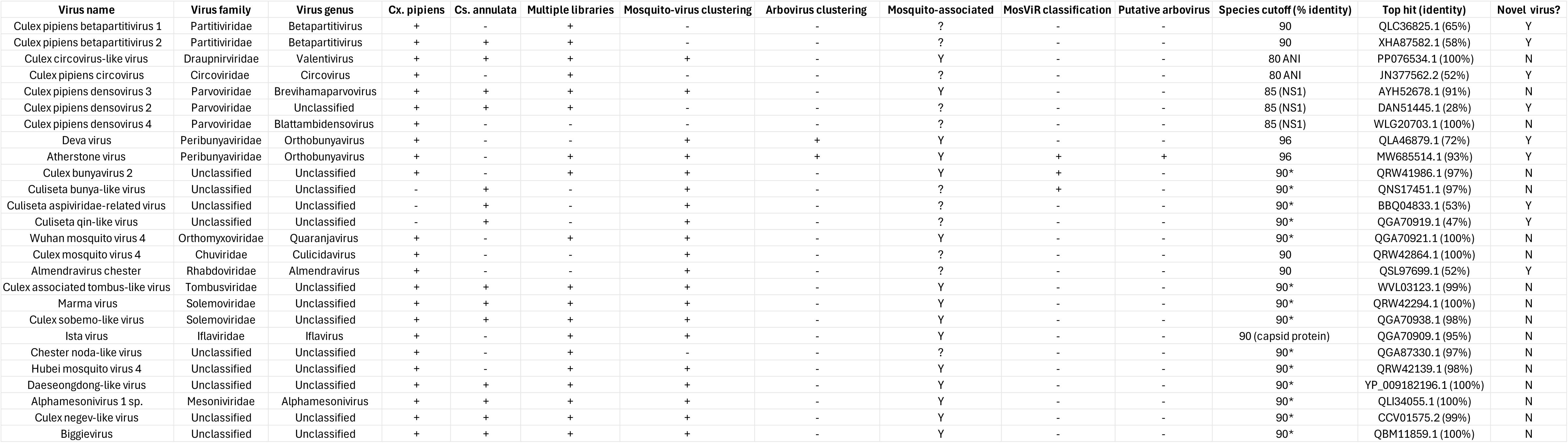
Summary of viruses identified in mosquito pools, including their taxonomic classification based on International Committee on Taxonomy of Viruses (ICTV) designation, host species presence, novelty status, and likelihood of being mosquito-associated or an arbovirus. Species demarcation is based on ICTV designation. In the absence of ICTV species demarcation, a threshold of <90% amino acid identity in the RNA-dependent RNA polymerase (RdRp) protein was applied relative to the closest NCBI match (*). Identity values refer to RdRp unless otherwise noted. ANI = indicates average nucleotide identity. A virus was considered a putative arbovirus if classified as such by the MosViR pipeline [29] as well as clustering phylogenetically with known arboviruses.

The number of viral reads detected in each pooled sample, as identified by Kraken2. NS1-NS47 correspond to female *Culex pipiens* s.l., NS48-NS50 correspond to female *Culiseta annulata*, and NSM1 and NSM3 correspond to male *Culex pipiens* s.l. Orange labels = Twycross zoo. Blue labels = Chester zoo. Background satellite imagery courtesy of OpenStreetMap contributors.

### Evolutionary history of UK mosquito viruses

Conserved proteins were used to reconstruct the evolutionary histories for each virus in order to assist in taxonomic assignment as well as inferences of host association and pathogenic potential. Most of the viruses (n = 15) clustered in clades where mosquitoes were prominent hosts or where arboviruses were present, and were also detected in multiple libraries, suggesting a likely mosquito association (Table 2). In contrast, 11 viruses clustered with those that had divergent or multiple hosts and/or were detected in only a single pooled sample, possibly indicating an alternative, non-mosquito source within the metagenome.

### Negative-sense RNA viruses

A total of 8 viruses were identified with a single-stranded negative-sense genome. These included members of the *Peribunyaviridae* (n=2), *Orthomyxoviridae* (n=1), *Chuviridae* (n=1) and *Rhabdoviridae* (n=1), with several others closely related to the *Qinviridae* (n=1), *Aspiviridae* (n=1) and *Phenuiviridae* (n=1) (Figure 2). The only viruses which had previously been described included Wuhan mosquito virus 4 (*Quaranjavirus*, *Orthomyxoviridae*) and *Culex* mosquito virus 4 (*Culicidavirus*, *Chuviridae*), as well as the bunya-like viruses (*Culiseta* bunya-like and *Culex* bunyavirus 2), which all form clades predominantly incorporating mosquito hosts. Novel viruses which were also likely mosquito viruses included *Culiseta* qin-like virus, which is part of a diverse clade solely comprising of viruses detected in mosquitoes (Figure 2) with its closest relative being Nackenback virus (QGA70919.1) with 47% amino acid identity. Similarly, a member of the *Rhabdoviridae* family placed in the Almendravirus genus which is almost exclusively found in mosquitoes and was related to Menghai virus (QSL97699.1; 52% amino acid identity). In contrast, an *Aspiviridae*-related virus with the closest relative being *Culex tritaeniorhynchus Aspiviridae-*related virus (53% amino acid identity; BBQ04833.1), formed a sister clade to exclusively fungal viruses in the proposed *Mycoaspiviridae* group [65] (Figure 2). Due to the lack of members in this new clade, it is unclear whether this is mosquito or fungal-related.

Notably, two novel species of *Orthobunyavirus* (*Peribunyaviridae*) were identified and coined Atherstone virus and Deva virus. The standard tripartite genome characteristic of the genus was detected for Atherstone virus with associated complete ORFs for each three segments (Figure 3). Its closest known relative is *Orthobunyavirus olifantsvleiense* (MW685514.1), a partial sequence detected in *Culex pipiens* from South Africa in 1963, sharing 93% amino acid identity in the RdRp protein. Supporting its classification as a putative arbovirus, Atherstone virus was predicted by the machine learning tool MosViR to have arboviral potential (M segment probability score = 0.64). Furthermore, an NSs open reading frame was identified in the S segment (Figure 3). It was located on the same strand as the N gene, initiated 73 nucleotides upstream of the N start codon, and encoded a 67-amino-acid peptide. Among all viral contigs assessed, only Atherstone virus, and the L segments of *Culex* bunyavirus 2 (probability score = 0.75), and *Culiseta* bunya-like virus (probability score = 0.62) met the threshold for arbovirus prediction via MosViR (Supplementary Table S2). However, as the pathogenic status of the bunya-like clade remains unclear, Atherstone virus was the only virus designated a putative arbovirus based on this study’s combined criteria.

In contrast, Deva virus was represented by a partial genome (full S segment ORF and partial L and M segments ORFs) and was most closely related to Weldona virus (QLA46879.1), previously detected in *Culicoides* biting midges from the USA in 1990 [66]. Deva virus lacked a detectable NSs ORF (Figure 3) and was not predicted by MosViR to have arboviral potential (Supplementary Table S2).

**Figure 2.**
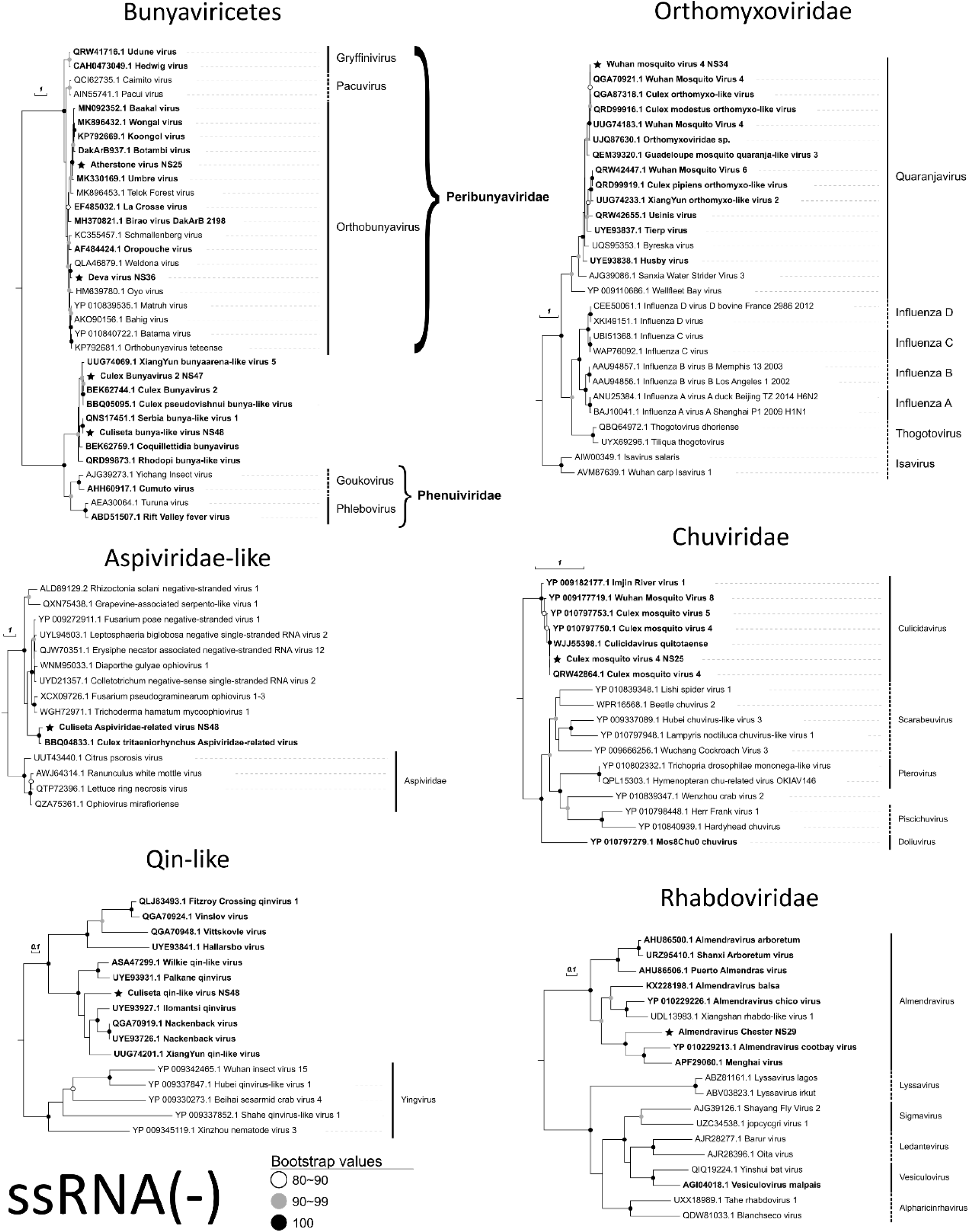
Phylogenetic analysis of negative-sense RNA viruses detected in this study (black stars) based on RNA-dependent RNA polymerase (RdRp) amino acid sequences. The analysis includes the closest matching sequences and representative members from ICTV-designated genera (solid or dashed lines) and families (brackets). Viruses identified in mosquitoes are highlighted in bold. Branch lengths reflect evolutionary divergence, measured by the number of amino acid substitutions per site.

**Figure 3.**
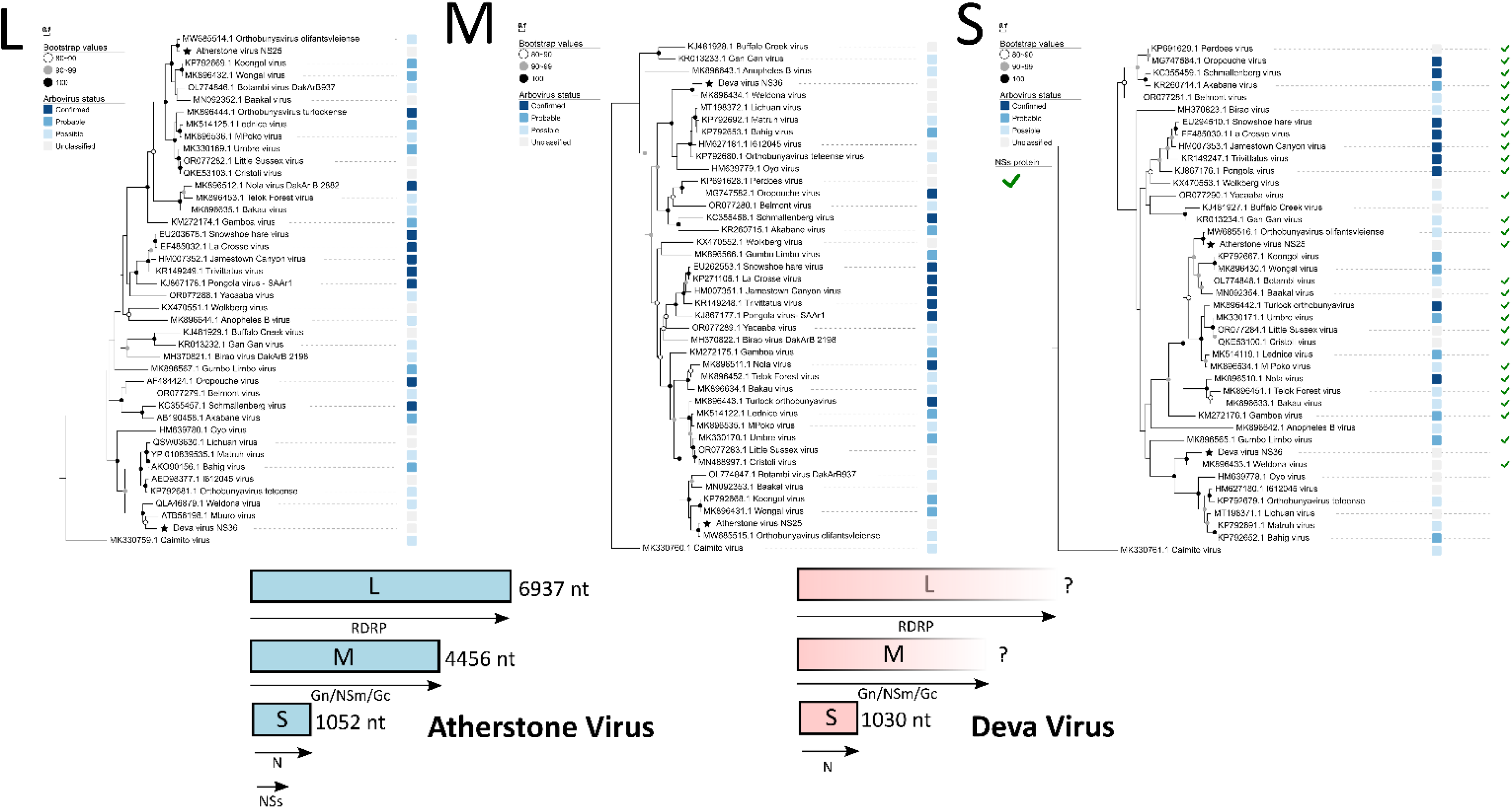
Phylogenetic analysis of the L (RNA-dependent RNA polymerase), M (Glycoprotein), and S (Nucleocapsid protein) segments of orthobunyaviruses. Black stars indicate viruses identified in this study. Annotations indicate the presence or absence of the non-structural protein NSs and the likelihood of arbovirus potential based on Centers for Disease Control and Prevention (CDC) Arbovirus Catalog classification [58]. Branch lengths reflect evolutionary divergence, measured by the number of amino acid substitutions per site. RdRp = RNA-dependent RNA polymerase; Gn = glycoprotein N; Gc = glycoprotein C; NSm = non-structural protein M; NSs = non-structural protein S; N = nucleocapsid.

### Positive-sense RNA viruses

Ten viruses were identified with a single-stranded positive-sense genome. These included, members of the *Solemoviridae* (n=2), *Tombusviridae* (n=1), *Mesoniviridae* (n=1), and *Iflaviridae* (n=1), with further undesignated viruses classified as Noda-like (n=1), Negev-like (n=2), and Daeseongdong-like viruses (n=1), as well as Hubei mosquito virus 4 (n=1) (Figure 4). Unlike negative-sense RNA viruses, no novel positive-sense viruses were found.

The detection of Alphamesonivirus 1, *Culex*-associated Tombus-like virus, and Daeseongdong-like virus in all samples, along with their phylogenetic signal with insect hosts, indicates a strong association with mosquitoes in the UK. Other viruses likely to be mosquito-specific include Ista virus, first discovered in *Culex pipiens* from Sweden [23], *Culex* Negev-like virus, Biggie virus, and Hubei mosquito virus 4 (Figure 4, Table 2). Additionally, Marma virus and *Culex* Sobemo-like virus, both members of the *Solemoviridae* family, clustered in an undesignated clade composed predominantly of mosquito hosts, despite other genera in the family being exclusively plant-associated. Similarly, a *Culex*-associated Tombus-like virus was detected within a clade composed exclusively of insect hosts, nested within the *Tombusviridae*, whose designated genera are also traditionally associated only with plant hosts.

The presence of these insect-associated clades within viral families previously thought to be plant-specific suggests that both *Solemoviridae* and *Tombusviridae* may have a broader host range than previously recognized. Similarly, viruses similar to Chester Noda-like virus appear to have a wide host range, encompassing both insects and vertebrates.

**Figure 4.**
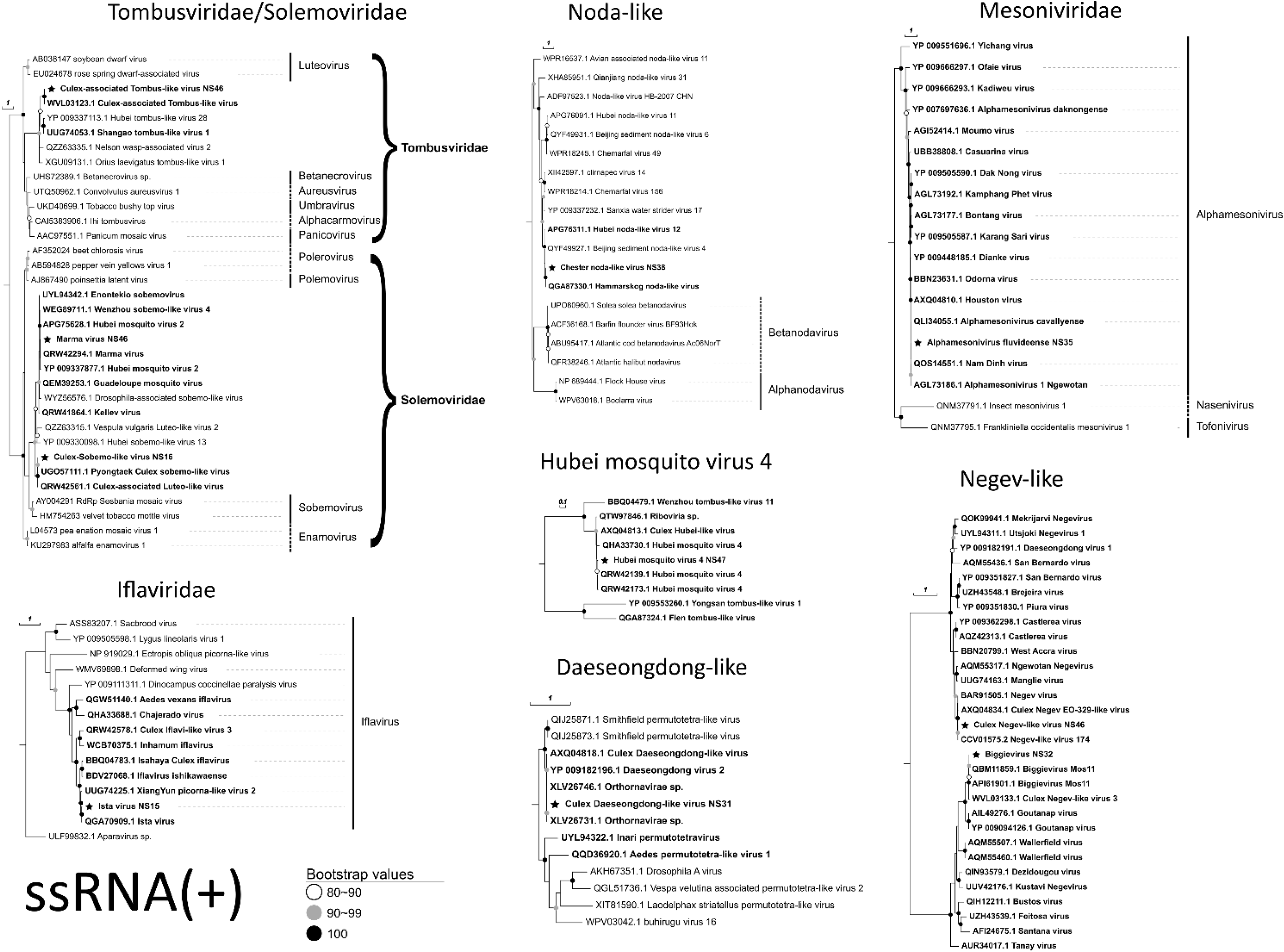
Phylogenetic analysis of positive-sense RNA viruses detected in this study (black stars) based on RNA-dependent RNA polymerase (RdRp) amino acid sequences. The analysis includes the closest matching sequences and representative members from ICTV-designated genera (solid or dashed lines) and families (brackets). Viruses identified in mosquitoes are highlighted in bold. Branch lengths reflect evolutionary divergence, measured by the number of amino acid substitutions per site.

### Double-stranded RNA viruses

Two novel viruses within *Betapartitivirus* (*Partitiviridae*) were detected (Figure 5). The first, *Culex pipiens* betapartitivirus 1, was identified exclusively in two sample pools from Chester Zoo. Phylogenetic analysis showed that it clusters closely with partitiviruses previously found in *Sarcosphaera coronaria* (QLC36825.1) and *Rosselina necatrix* fungi (BBU59838.1). In contrast, *Culex pipiens* betapartitivirus 2 was found in a majority of pools (n=27) with its closest relative being a virus from the freshwater crayfish *Procambarus clarkii* (XHA87582.1).

### Single-stranded DNA viruses

Five distinct ssDNA viruses were identified falling into the *Parvoviridae*, *Circoviridae* and *Draupnirviridae* families (Figure 6). Evolutionary reconstruction of the replication-associated protein (Rep) in *Circoviridae* and *Draupnirviridae* placed the two viruses within the genera Circovirus (*Culex pipiens* circovirus) and Valentivirus (*Culex* circovirus-like virus), respectively. The latter genus, which has primarily been described in bats and *Culex* mosquitoes [67], includes a virus nearly identical to the circovirus-like virus detected in this study (NC_040576.1). In contrast, the detected member of the *Circovirus* genus has a closest known relative from bats (52% amino acid identity; JN377562.2).

Of the three *Parvoviridae* members (*Culex pipiens* densoviruses 2, 3, and 4), only *Culex pipiens* densovirus 2 formed a distinct clade separate from designated genera. According to ICTV criteria, genus demarcation requires less than 85% amino acid identity in the NS1 protein between strains, and *Culex pipiens* densovirus 2 shared only 28% identity with its closest match, a *Parvoviridae* sp. from a human (DAN51445.1). In contrast, *Culex pipiens* densovirus 3 grouped within the *Brevihamaparvirus* genus, while *Culex pipiens* densovirus 4 (100% identity to *Zophobas morio* black wasting virus; WLG20703.1) clustered within *Blattambidensovirus*.

**Figure 5.**
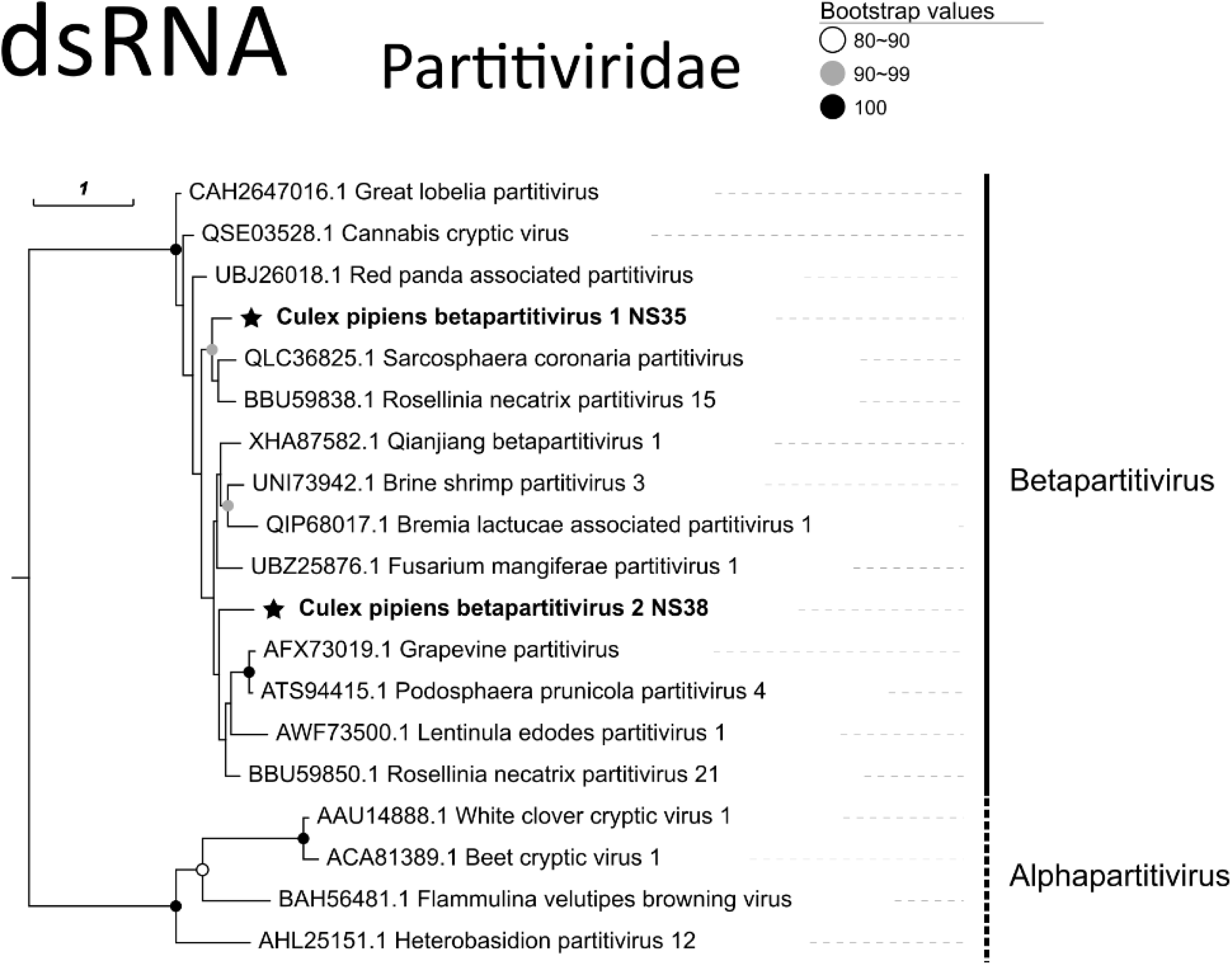
Phylogenetic analysis of double-stranded RNA (dsRNA) viruses detected in this study (black stars). The tree is reconstructed from RNA-dependent RNA polymerase (RdRp) amino-acid sequences and includes the closest matches from public databases as well as representative members of the ICTV-designated Partitiviridae family, as well as betapartitivirus and alphapartitivirus genera. Mosquito-derived viruses are highlighted in bold. Branch lengths are proportional to the number of amino-acid substitutions per site.

**Figure 6.**
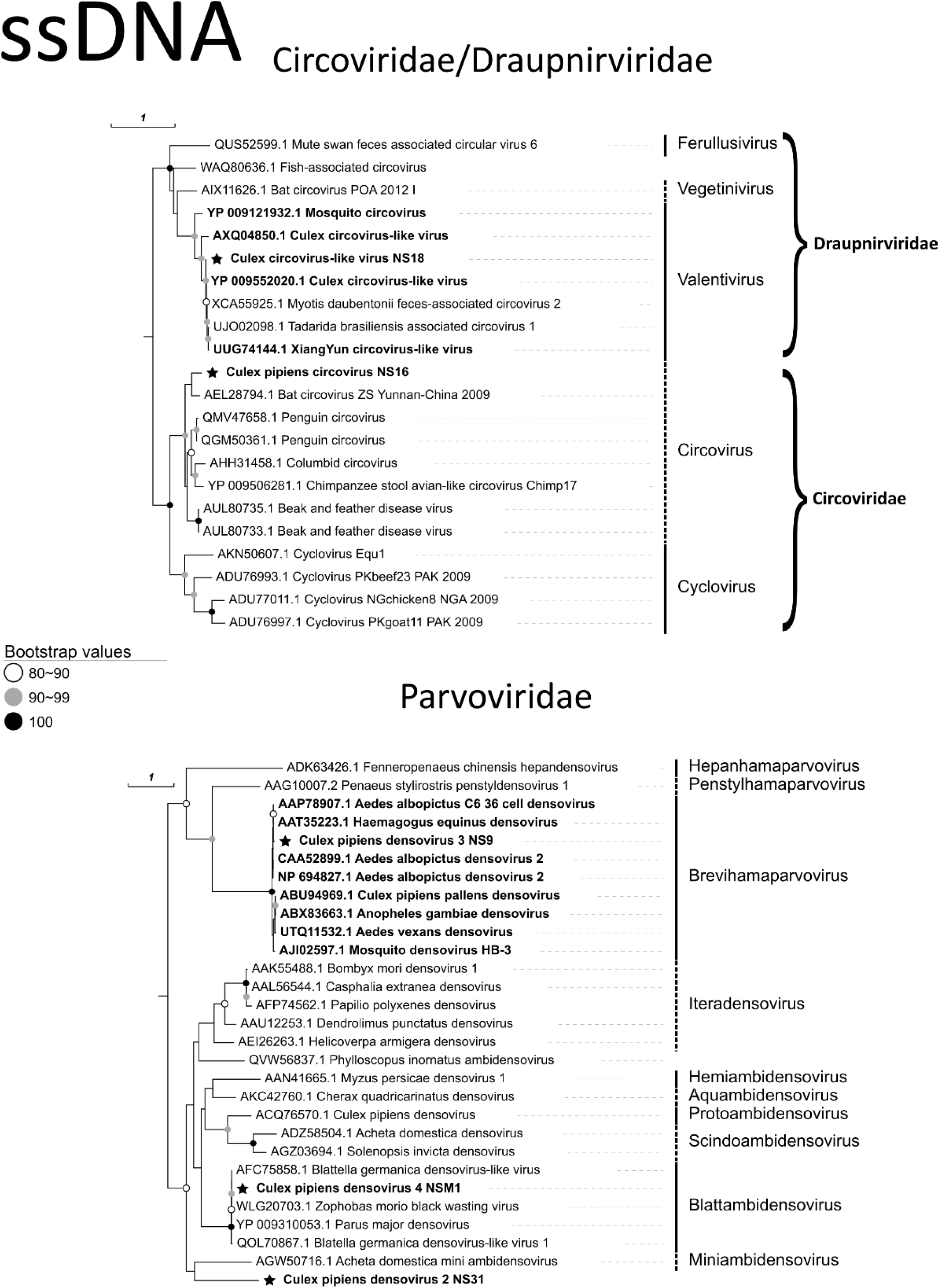
Phylogenetic analysis of single-stranded DNA (ssDNA) viruses detected in this study (black stars). For Circoviridae and Draupnirviridae, replication-associated protein sequences were used; for Parvoviridae, NS1 protein sequences were used. The tree includes the nearest database matches and representative ICTV genera (solid lines) and families (brackets). Mosquito-derived ssDNA viruses are shown in bold. Branch lengths reflect estimated amino-acid substitutions per site.

### Patterns of Virus Detection in Mosquito Samples

For each library, the number of virus species varied from 5 to 16 (Figure 7). Furthermore, five viruses were detected consistently in each pool from both sites. These included *Culex* densovirus 3, *Culex* Circovirus-like virus, Alphamesonivirus 1, as well as Tomus-like and Daeseongdong-like viruses. When comparing the viromes of *Culex* and *Culiseta* pools, both shared further viruses including *Culex* densovirus 2, Marma virus, Biggievirus, *Culex* pipiens betapartitivirus 2, as well as Sobemo-like and Negev-like viruses. Their widespread presence suggests efficient transmission through high-fidelity horizontal and/or vertical transmission routes.

**Figure 7.**
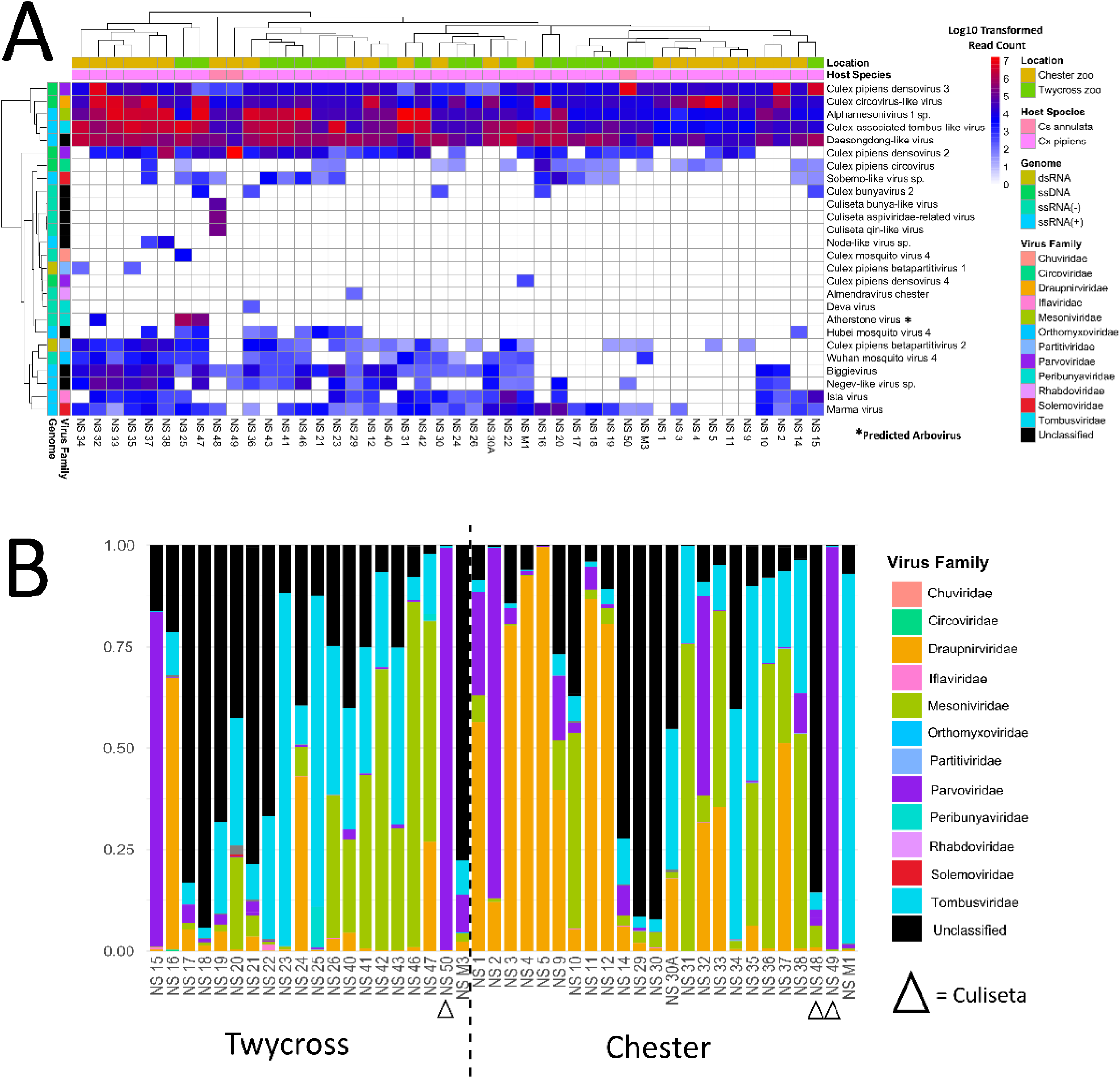
**(A)** Heatmap showing viruses detected in *Culex pipiens* and *Culiseta annulata* pools from Chester and Twycross zoos (2021–2022). A virus was classified as a putative arbovirus (asterisk) based on predictions from the machine-learning tool MosViR [29] and its phylogenetic clustering with known pathogenic clades. **(B)** Relative abundance of viral families in each pool. For both panels, viral families were designated based on ICTV criteria.

Unique viruses to *Culiseta* included a bunya-like virus, an *Aspiviridae*-like virus and a qin-like virus, which were all present in 1 of 3 *Culiseta* pools but absent in the 41 *Culex* pools (Figures 7 and 8). Notably, while most viruses (n=17) were detected at both Chester Zoo and Twycross Zoo, some were unique to specific mosquito species at each site. A small number were found exclusively in *Culex pipiens* (n=5) and *Culiseta annulata* (n=3) at Chester Zoo, while only one virus (n=1) was unique to *Culex pipiens* at Twycross Zoo (Figure 8), indicating limited geographic restriction of viral species.

**Figure 8.**
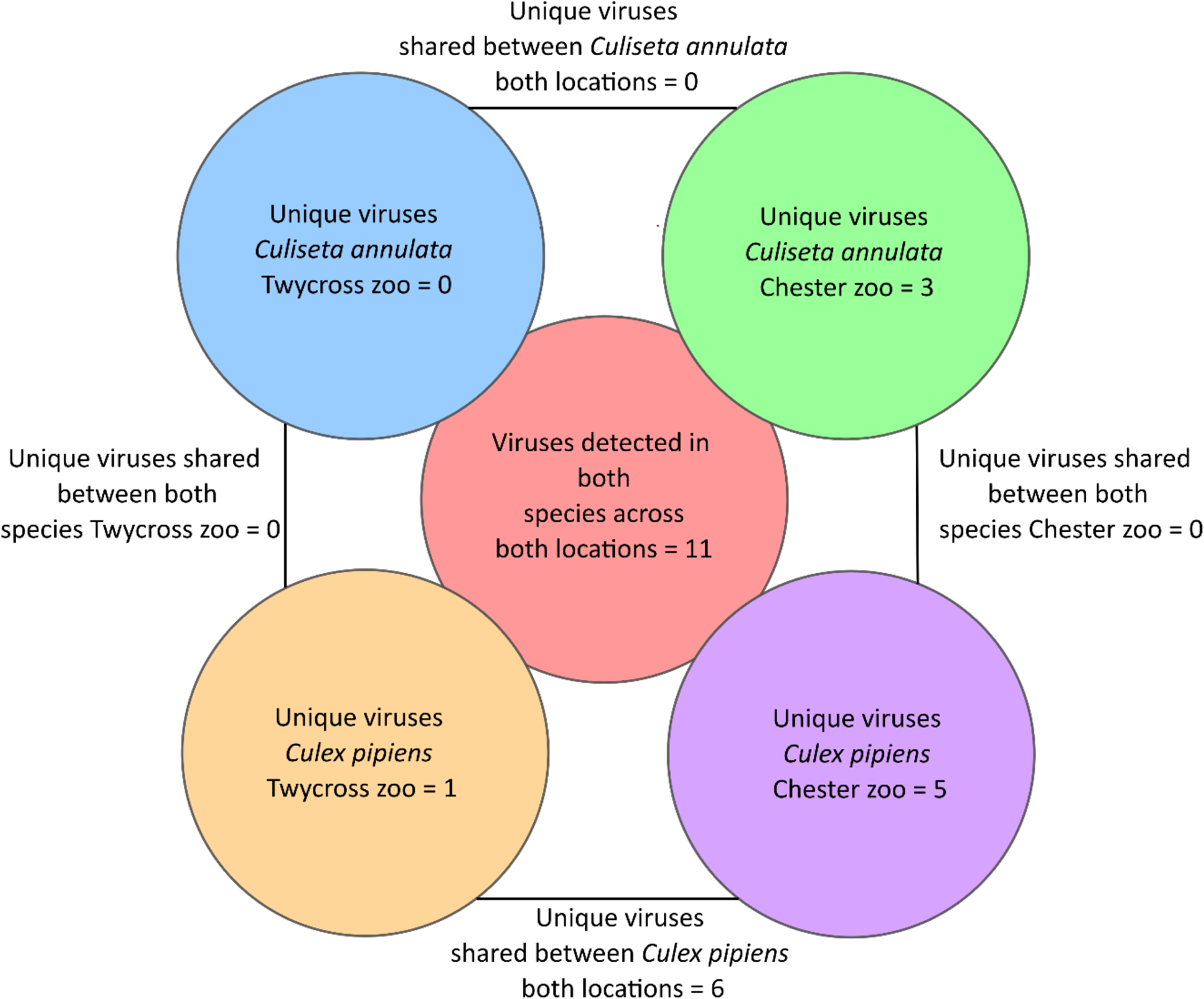
Venn diagram showing the number of shared and unique viruses between both species and both zoos.

Among viruses detected in multiple pooled samples, 13 produced multiple reference-based assemblies (using the *de novo* assembled reference) that met the criteria of 10× sequencing depth and 90% genome breadth coverage (Figure 9). All of these exhibited high ANI values, with an average of 99.68 ± 0.26 for ssDNA viruses, 99.89 ± 0.11 for dsRNA viruses, 99.26 ± 0.53 for ssRNA(+) viruses, and 98.84 ± 0.45 for ssRNA(-) viruses. These consistently high ANI values indicate a relatively low level of genetic divergence, while the observed variation suggests genuine viral diversity rather than cross-contamination.

**Figure 9.**
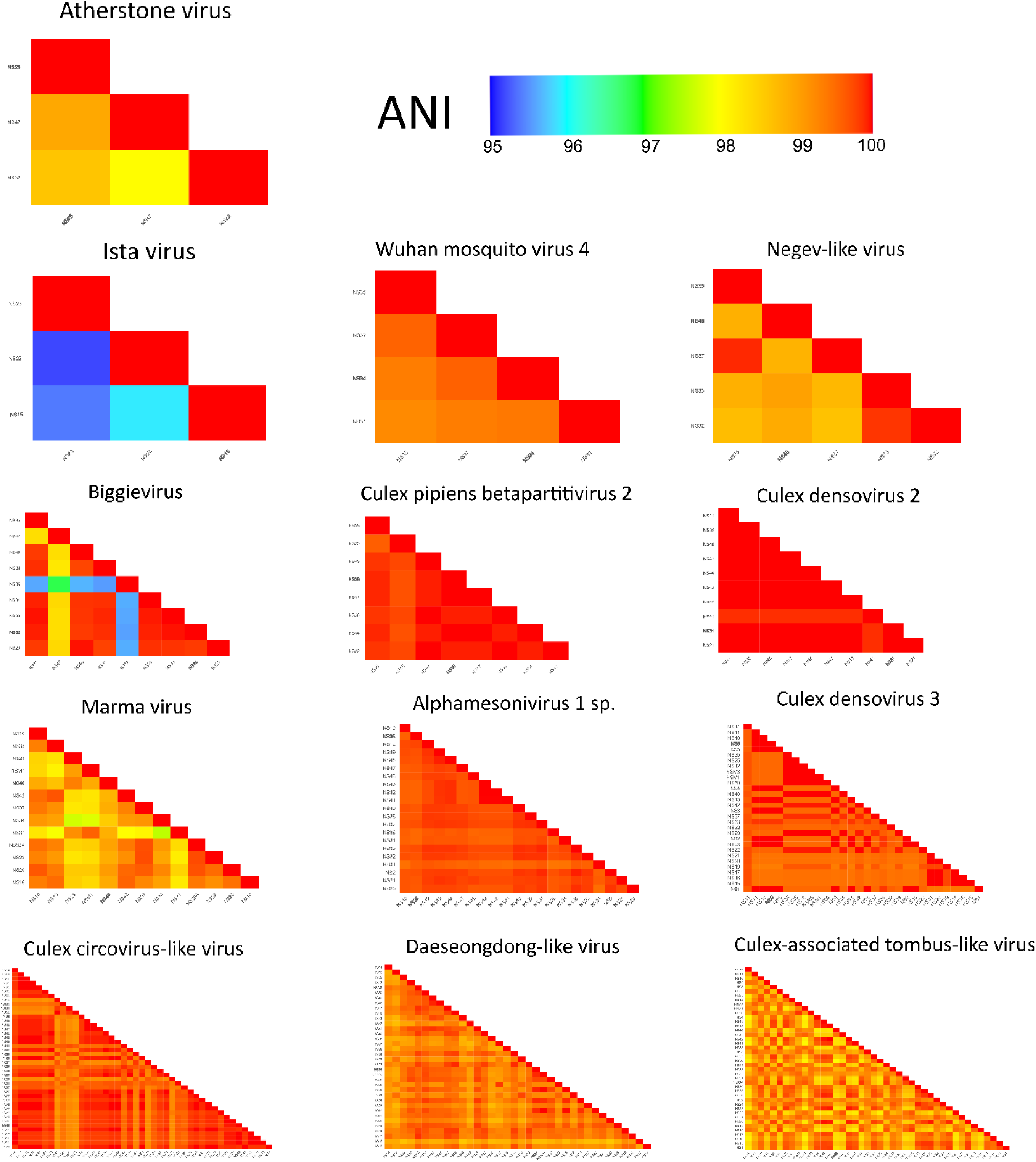
Heatmap showing the genomic similarity of viral genomes across pooled samples, based on Average Nucleotide Identity (ANI). Bold sample IDs refer to reference genomes from *de novo* assemblies.

## Discussion

In this study, 26 viruses including 9 novel species were detected from 2 UK zoos, of which some are candidates warranting further investigation for their potential public and animal health implications. Most strikingly we detected two viruses within *Orthobunyavirus*, Atherstone and Deva viruses, marking the first documented mosquito-associated orthobunyaviruses in the UK. While much of the UK’s mosquito-borne disease research has focused on USUV [2,3,41], which is already established, and WNV which is anticipated to emerge in the near future [68], little attention has been paid to other potentially pathogenic viruses that may already be circulating in native mosquito populations. The presence of these orthobunyaviruses highlights a potential gap in current surveillance efforts, as they may represent previously unrecognised, yet existing, threats. Orthobunyaviruses are a diverse group within the *Peribunyaviridae* family, many of which have significant public and veterinary health implications [69]. Notably, this group contains several known arboviruses such as Schmallenberg virus [8], which causes congenital abnormalities and abortion in ruminants, as well as the zoonotic Oropouche virus which has recently expanded its known range in Latin America [70].

More recently, deaths due to orthobunyaviruses have been documented in Europe, including fatal encephalitis cases in France caused by Cristoli virus [71] and Umbre virus [72]. These examples highlight that orthobunyaviruses are not limited to tropical regions and can pose significant risks to public health in temperate zones. In this context, the discovery of Atherstone virus, detected at both sampled sites, is notable. It is predicted by MosViR to have arboviral potential, and critically, encodes a NSs protein. The NSs protein, a known virulence factor involved in immune suppression and maintenance of infection in vertebrates [56,57], further strengthens the case for Atherstone virus as a putative arbovirus of concern. In contrast, Deva virus, lacked both an NSs ORF and arbovirus classification by MosViR, but still warrants attention given its placement within the *Orthobunyavirus* genus.

While these viruses were identified in mosquito populations collected at two zoos, no evidence currently links their presence to disease in animals or humans at these sites. However, given the proximity to both exotic and native vertebrate species, these findings support the value of continued metagenomic surveillance in sentinel sites. Zoological collections, in particular, offer an opportunity to detect potential spillover or introduction events at an early stage [73]. Future studies should include vertebrate serosurveys to assess prior exposure in animal or human populations, and *in vitro* studies to confirm replication competence in vertebrate cells. If vertebrate infection is demonstrated, evaluation of vector competence and pathogenicity in animal models would be warranted.

Alongside the detection of orthobunyaviruses, the consistent presence of an Alphamesonivirus in all libraries also raises potential concerns for animal health. This virus, previously believed to be incapable of replicating in mammalian cell lines [74], was recently identified in the lymph nodes and lung tissue of two horses suffering from respiratory syndromes [75]. Its uncertain classification as an insect-specific virus, coupled with its widespread presence in the samples analysed in this study, suggests that it may be a potential pathogen warranting consideration in cases of unexplained animal illnesses.

Another notable observation was the high abundance of *Culex* circo-like virus, the most dominant taxon in terms of viral reads, which belongs to the recently designated family *Draupnirviridae* [67]. The prototypical member of this group was first discovered in *Aedes vexans* [76] and classified under a new ICTV-designated genus, *Valentivirus* (also known as *Krikovirus* [[76]]). A recent study investigating the evolutionary history of this lineage has revealed endogenous viral elements (EVEs) integrated into the genomes of reptiles [67]. These elements, along with frequent infections observed in mosquitoes and bats, suggest that vertebrate infections may have been historically vector-mediated. Although the *Krikovirus* identified here did not meet this study’s criteria for classification as a putative arbovirus, its high abundance and known associations with mosquitoes and bats warrant continued surveillance to clarify its ecological role and potential vertebrate tropism.

To contextualise the broader origins and host associations of the detected viruses, phylogenetic analysis revealed that 15 likely originated from mosquitoes, as they clustered within clades associated with mosquito hosts and recurred across samples. Additionally, the observed overlap in virome composition between *Culex pipiens* and *Culiseta annulata* suggests that their shared ecological niches contribute to similar viral profiles. Notably, five viruses were consistently detected across all sample pools and both zoo sites, indicating their widespread presence in UK mosquito populations. Their detection in both male and female *Culex pipiens* further implies stable maintenance through high-fidelity horizontal and/or vertical transmission, rather than transient acquisition via blood feeding. While a few viruses were unique to each zoo, the consistent presence of several viruses across species, samples, and locations suggests that host species and transmission dynamics could be more influential than geography in shaping virome composition.

Taken together, these patterns align with previous studies suggesting the existence of a conserved core virome within and between mosquito species [23,77–79], potentially reflecting similar dynamics within UK populations. However, other large-scale studies—such as those examining mosquito viromes in China [80] and Belgium [25]—challenge this notion, indicating that viral communities are highly dynamic and shaped by geographical and seasonal variations. For this reason, it remains uncertain whether the mosquito-associated viruses in this study reflect stable, long-term associations maintained through transmission, or are instead shaped by geographical and environmental factors. To resolve this, single-mosquito metagenomics across diverse UK regions and timepoints could help disentangle these drivers.

A group of note detected in this study were members of the *Parvoviridae* family [38,81,82]. Viruses identified were classified as mosquito densoviruses (MDVs) within the genera *Brevihamaparvovirus* (*Culex* densovirus 3) and *Blattambidensovirus* (*Culex* densovirus 4), as well as one undesignated species (*Culex* densovirus 2). MDVs do not typically cause cytopathic effects in cell lines [83], but they have sometimes been shown to be pathogenic to mosquito larvae through oral infection [84]. Due to their larvicidal properties, host specificity, and multiple transmission routes, MDVs are considered promising candidates for mosquito population control, either through direct infection or viral paratransgenesis [38,84,85].

The widespread presence of *Culex* densovirus 3 (in all samples) and *Culex* densovirus 2 (in most samples) suggests that these viruses are well established in UK mosquito populations. While their abundance could indicate low pathogenicity, it is also possible that they persist through vertical transmission or sublethal infections, maintaining mosquito populations despite potential fitness costs. In contrast, *Culex* densovirus 4, which was found in only one pool is genetically identical to *Zophobas morio* black wasting virus, a pathogen responsible for widespread and rapid mortality of larval superworms (*Zophobas morio*) [86]. Its limited detection in this study may reflect low prevalence rather than low virulence, and further investigation is needed to assess its potential as a mosquito population suppression agent.

## Conclusion

This study demonstrates the value of metagenomic surveillance in improving our understanding of the diversity of mosquito-associated viruses in the UK, including the detection of novel species and arbovirus candidates. These findings provide important baseline data on the UK mosquito virome and highlight previously unrecognised viral taxa that may have implications for animal and public health. Continued monitoring using these approaches will be essential for understanding virus transmission dynamics, identifying emerging pathogens, and informing targeted vector control and surveillance strategies.

## Data availability

Virus sequences and raw reads are available at the European Nucleotide Archive under project accession number PRJEB87852.

## Author contributions

NS, ACD, MB, MSCB and GLH secured funding for this project. JP, NS and ACD contributed to the conceptual development of the project. NS, PD and JL assisted in carrying out fieldwork. NS and JP conducted laboratory work. JP and ECO carried out bioinformatic analysis. JP, NS, ECO, AK, GLH, MSCB, MB and ACD interpreted data. JP produced figures and drafts of the manuscript. All authors assisted in critical revision of the manuscript.

## Acknowledgements

The authors would like to thank Lindsay Eckley and Helena Turner at Chester Zoo for their assistance in specimen collection, and the Birds, Primates, and Veterinary teams at Twycross Zoo, as well as Lisa Gillespie for their support during fieldwork. We also acknowledge the support of Dr. Richard Gregory and the Centre for Genomic Research (CGR) at the University of Liverpool for providing access to computational resources used in this research, as well as for the use of CGR’s sequencing facilities.

